# A national professional development program fills mentoring gaps for postdoctoral researchers

**DOI:** 10.1101/2022.09.26.509546

**Authors:** Ting Sun, Denise Drane, Richard McGee, Henry Campa, Bennett B Goldberg, Sarah Chobot Hokanson

## Abstract

*The Postdoc Academy: Succeeding as a Postdoc* was designed to build postdocs’ skills in career transition, career planning, collaborative research, resilience, and self-reflection. This study examined self-reported changes in five skills as learners progressed through the course. Data were collected from participants who responded to both pre- and post-surveys and engaged with the course learning activities. Results from repeated measures multivariate analysis of variance revealed that all of the self-reported perceptions of skills improved significantly upon completion of the course. Hierarchical regressions revealed that underrepresented minority learners had greater gains in their development of skills in career planning, resilience, and self-reflection. Qualitative analysis of learners’ responses to learning activities found that postdocs perceived networking and mentor support as contributing factors to their skill advancement while tensions among multiple obligations and concerns of uncertainties were significant challenges to applying those skills.

## Introduction

Postdoctoral scholars (hereafter “postdocs”) have long been acknowledged as important contributors to scientific and economic advancement [1]. To achieve research independence and professional readiness, postdocs benefit from effective career and professional development [2]. Professional development has a positive impact on postdoctoral experiences and future career success [3], as those who receive more professional development tend to have greater overall life and job satisfaction, fewer conflicts with their advisers, and more publications and grant submissions [3, 4]. Hokanson et al. found that nine hours of synchronous, online development workshops were important to self-reflective practices and skill-building processes for graduate students and postdocs [5]. A meta-analytic study also demonstrated that professional development was a significant predictor of upward career mobility and was associated with higher salary, promotion and career satisfaction [6]. These results have sparked an emerging interest in professional development and have motivated institutions’ postdoc offices and funding agencies to design and implement career and professional development programs for postdocs. A small sampling of these include: Preparing Future Faculty Program support by the Council of Graduate Schools and the American Association of Colleges and Universities; the Burroughs Wellcome Fund Career Guidance for Training award; NIH National Institute of General Medical Science, Innovative Program to Enhance Research Training grant; and, most recently, the NIH Broadening Experiences in Scientific Training (BEST) program [7], which aimed to provide career training opportunities for biomedical doctoral and postdoctoral researchers.

### Statement of the problem

Numerous challenges exist with current professional development opportunities for postdocs offered at academic institutions and/or through professional societies. Many programs focus on academic careers (e.g., The Preparing Future Faculty Program) and development for non-academic career paths can be limited [8]. This, despite the fact that only 15% of postdocs secure tenure-track faculty positions [2, 9]. Second, some professional development opportunities cannot reach their target audiences because their offerings conflict with postdocs’ research and personal schedules [10]. Lastly, some relatively large-scale professional development opportunities focus exclusively on biomedical disciplines largely ignoring other disciplines (e.g., BEST) [7]. We have designed, implemented, and evaluated a professional development program that offers a broad focus on skills applicable to academic and non-academic careers, flexible delivery and engagement, and covers a diversity of career opportunities for postdocs across disciplines.

Mentoring plays an essential role in supporting postdocs in their professional development engagement [11]. Yet some mentors are either unaware of professional development opportunities (see above) or may not see the benefit to their postdoc’s research and career success [12]. Some faculty mentors also have limited time, and limited experience and access to resources to advise their mentee postdocs in non-academic career pathways [13]. From either advisors or institutional programs, some postdocs are not provided with consistent and sufficient resources or opportunities for professional development [13]. This lack of resources has resulted in inadequate mentoring support and postdocs have difficulty in balancing their research and professional development engagement [12]. The situation is even worse with underrepresented minorities (URM) and females [14, 15]. URM refers to individuals from racial and ethnic groups such as Black or African American, Hispanic or Latino, American Indian or Alaska Native, Native Hawaiian, and other Pacific Islander [16]. URM postdocs and early career faculty were found to have less mentoring as compared to their majority counterparts [17]. Additionally, due to marginalities derived from their ethnicity and gender, URM postdocs have more challenges in accessing and navigating professional development opportunities and resources [12, 18]. Developing and implementing an open access, inclusive and structured professional development program could help meet postdoc mentoring needs or supplement interpersonal and institutional mentoring, which can contribute to success for diverse postdocs.

### The Postdoc Academy program

The Postdoc Academy is a comprehensive professional development program for postdocs everywhere, designed to reach those in both research-intensive and well-resourced institutions, as well as those who are isolated with limited local networking or professional development opportunities. The Postdoc Academy was motivated by the national reports that described the needs of postdocs [19], the recognition that they are an oft-ignored segment of the academic pathway, and the strong positive impact of professional development on postdoc career success [3]. The Postdoc Academy, which consists of two online asynchronous courses that also have optional in-person synchronous meetings, aims to build skills aligned with the National Postdoctoral Association core competencies [20], and supports career advancement throughout the postdoc pathway. As a blended learning program, it offers multiple learning modalities to reach diverse audiences. This program developed and implemented two Massive Open Online Courses (MOOCs) offered on the edX platform, *The Postdoc Academy: Succeeding as a Postdoc* and *The Postdoc Academy: Building Skills for a Successful Career*. This paper examines the first course that was launched, *The Postdoc Academy: Succeeding as a Postdoc*.

*Succeeding as a Postdoc* consists of four modules, *Finding Success as a Postdoc, Building an Actionable Career Plan, Developing Resilience*, and *Working Effectively in an Intercultural Environment*, with the aim to build learners’ skills in career transition, career planning, collaborative research, resilience and self-reflection. Learners can participate in the course by watching course videos or reading video transcripts, participating in discussions, completing individual reflections, and engaging with interactive learning activities (Fig 1). The course was offered five times from January 2020 through January 2022. Each offering lasted six weeks, with one being offered before the COVID-19 pandemic and four being offered after the pandemic.

**Fig 1.**
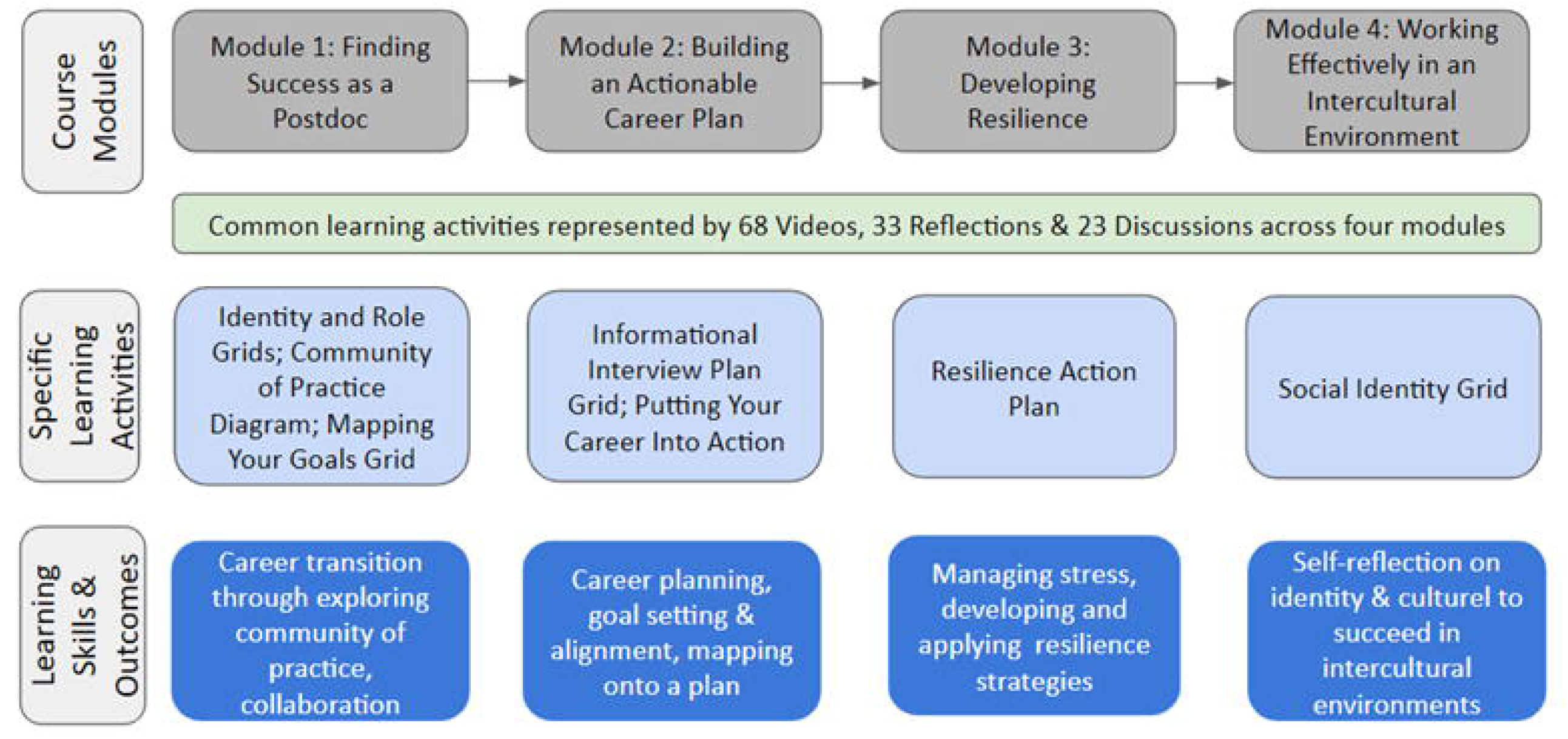
Modules, Learning Activities, and Learning Skills and Outcomes of *The Postdoc Academy: Succeeding as a Postdoc* from February 2020 through January 2022.

### Effects of skill building on postdoctoral success

Previous literature suggested that skills in career transition, career planning, collaborative research, resilience and self-reflection play a vital role in postdoc professional and academic success [6, 21]. Career planning and transition are critical for achieving short- and long-term goals. Ng et al. [6] conducted a meta-analysis on the predictors of career success and found that career planning was positively and significantly correlated with objective career success (i.e., greater salary, *r* = .11, *p* < 0.05) and subjective career success (i.e., career satisfaction, *r* = .33, *p* < 0.05). Resilience is defined as “positive adaptation in the face of risk or adversity” or “capability of a dynamic system to withstand or recover from disturbance” (p. 17) [22] and also plays a vital role in postdoc success. Gloria and Steinhardt examined the relationships among positive emotions, coping, resilience and mental health in a sample of 200 postdocs and found that resilience was positively correlated with positive emotions and adaptive coping and negatively correlated with maladaptive coping, stress, trait anxiety, and depressive symptoms [21]. Collaboration is another essential professional skill used in numerous disciplines to tackle complex problems in research or management and to develop creative works [23]. The complexity of many problems often requires specialized knowledge from multiple disciplines necessitating collaboration among scientists to bring together their expertise and skills [24]. Scaffidi and Berman examined factors influencing the experiences and research productivity in a sample of 204 postdocs and found that research collaboration had a positive effect on research output [23].

Previous research has suggested a positive correlation between career transition, planning, collaborative research, resilience and self-reflection and postdoc success. Since these skills are central to the learning outcomes of *Succeeding as a Postdoc*, it is important to examine correlations in our learners associated with the development of these skills. It is also important to examine how different demographic groups navigate professional development opportunities, since existing literature indicates that URM and female postdocs had unique challenges navigating professional development opportunities [12]. We sought to examine the learner perception changes in the five skills as learners progressed through the course. The research questions that guided our study are:

1. To what extent did learners’ self-reported skills (career transition, career planning, collaborative research, resilience, and self-reflection) change after they participated in *The Postdoc Academy: Succeeding as a Postdoc*?
2. What are the differences in the changes of learners’ self-reported skills as a function of learners’ demographic characteristics (i.e., gender, ethnicity, discipline, and country of origin)?
3. What were learners’ reflections on skill development as described in their learning experiences as they engaged with *The Postdoc Academy: Succeeding as a Postdoc*?

## Materials and methods

### Study design

This study used a mixed methods design with quantitative and qualitative methods. A single-group, pretest-posttest design was used to measure self-reported learning gains and behavioral intentions. Although not the strongest experimental design, the single-group design is widely used in education situations where having more than one form of instruction simultaneously is not feasible [25]. Qualitative analysis of learners’ direct work products (course learning activities) was incorporated to elicit in-depth answers to the first two research questions and to enrich our understanding. A detailed description of the course learning activities can be found in the Data Sources section.

### Participants

Study participants were those who responded to both pre- and post-course surveys. The number of survey respondents, engaged learners, registrants, and completers for each of the *Succeeding as a Postdoc* offerings are presented in Table 1. A total of 5581 learners enrolled in *Succeeding as a Postdoc* and 473 completed the course. The total numbers of survey respondents were 1334 and 376 for the pre- and post-course surveys, respectively. The current study used the sample who responded to both the pre- and post-course surveys. Twenty-three participants who did not provide consent to use their survey responses for research were removed from the study, yielding 215 participants. A subset of the 215 participants ranging from 166 to 209 who interacted with *Succeeding as a Postdoc* course learning activities was also created for the learning activities analysis.

**Table 1.**
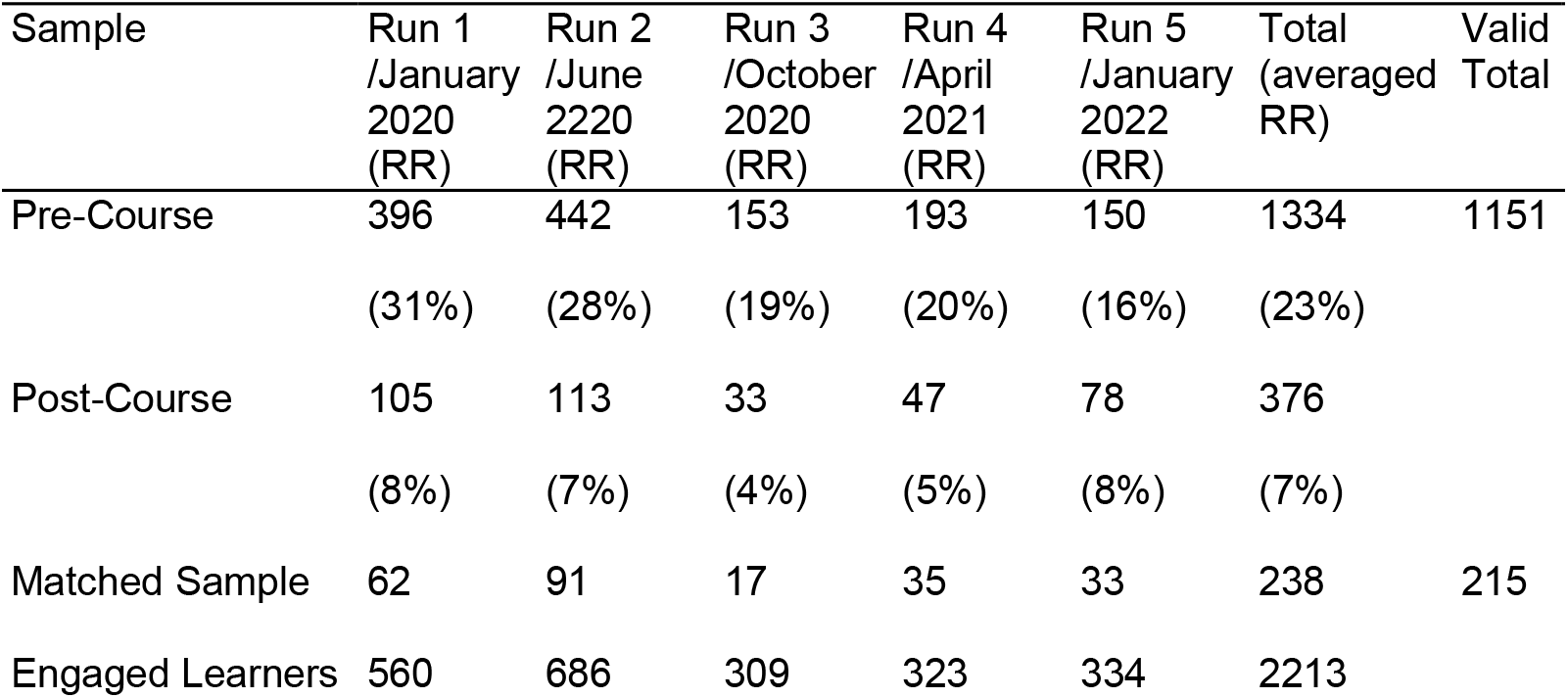
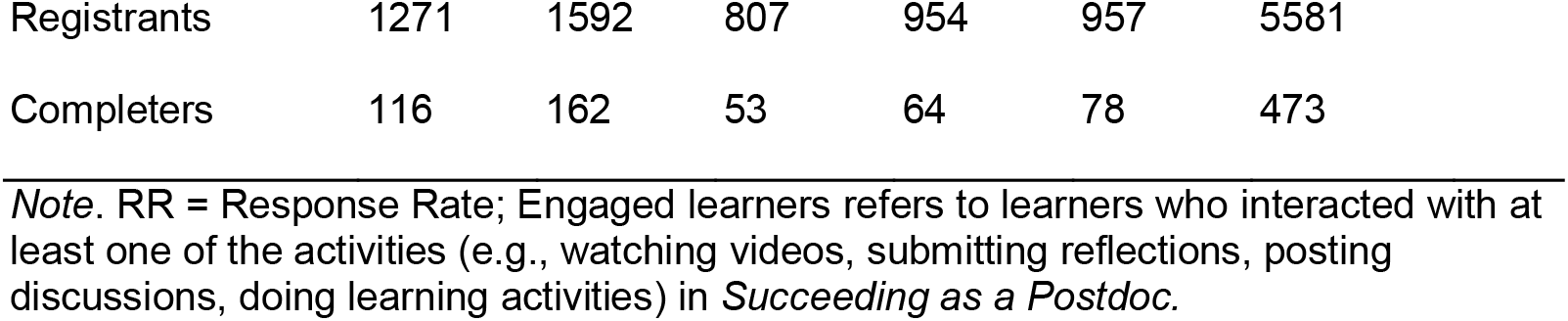
Number of Survey Respondents, Engaged Learners, Registrants, and Completers by Course Run from *The Postdoc Academy: Succeeding as a Postdoc* (from February 2020 through January 2022)

| Sample | Run 1<br>/January<br>2020<br>(RR) | Run 2<br>/June<br>2020<br>(RR) | Run 3<br>/October<br>2020<br>(RR) | Run 4<br>/April<br>2021<br>(RR) | Run 5<br>/January<br>2022<br>(RR) | Total<br>(averaged<br>RR) | Valid<br>Total |
| --- | --- | --- | --- | --- | --- | --- | --- |
| Pre-Course | 396<br>(31%) | 442<br>(28%) | 153<br>(19%) | 193<br>(20%) | 150<br>(16%) | 1334<br>(23%) | 1151 |
| Post-Course | 105<br>(8%) | 113<br>(7%) | 33<br>(4%) | 47<br>(5%) | 78<br>(8%) | 376<br>(7%) |  |
| Matched Sample | 62 | 91 | 17 | 35 | 33 | 238 | 215 |
| Engaged Learners | 560 | 686 | 309 | 323 | 334 | 2213 |  |

|  |  |  |  |  |  |  |
| --- | --- | --- | --- | --- | --- | --- |
| Registrants | 1271 | 1592 | 807 | 954 | 957 | 5581 |
| Completers | 116 | 162 | 53 | 64 | 78 | 473 |

*Note.* RR = Response Rate; Engaged learners refers to learners who interacted with at least one of the activities (e.g., watching videos, submitting reflections, posting discussions, doing learning activities) in *Succeeding as a Postdoc*.

The demographic information for the sample who responded to both surveys is shown in <u>S1 Table</u>. The sample is predominantly postdocs (*n* = 178, 83%) and females (*n* = 146, 68%). White/Caucasian participants made up 42% of the sample, followed by Asian or Asian American (19%), Hispanic or Latino/Latina/Latinx (12%) and smaller numbers of other racial/ethnic groups. Approximately half of the learners (56%) worked in biological/medical sciences, and learners working in physical/engineering/computer sciences and humanities/social sciences represented 14% and 15% of the sample, respectively. Learners from 37 countries of origin were included in the sample, with the United States accounting for the largest proportion (43%). The demographic composition of pre-course survey respondents and the national postdoc population are also presented in <u>S1 Table</u> to demonstrate the representativeness of the sample used in the study. The demographic composition of the sample is similar to those who responded to the pre-course survey. However, our sample had more females, more URM learners, and learners working in humanities/social sciences compared with the national postdoc survey respondents [26].

## Data Sources

### Surveys

Survey instruments used in the current study include pre- and post-course surveys. Surveys were designed by embedded evaluators and course designers based on the theory of change for each of the four course modules, which connected course pedagogical design, learning activities and short-term outcomes [27]. The pre-course survey had five items asking participants to rate (using a five-point Likert scale; 1 = Not proficient to 5 = Extremely proficient) their self-perceptions of their competency in making career transitions skillfully, planning career, developing collaborative research relationships, rebounding from setbacks or challenges, and engaging in self-reflection, which are central to the learning goals and theory of change of the modules in *Succeeding as a Postdoc*. The pre-course survey also had items collecting participants’ demographic information including gender, ethnicity, discipline, and country of origin. The post-course survey included the same five items assessing participants’ perceptions of their skills upon completion of *Succeeding as a Postdoc*. The post-course survey also had items evaluating the quality of each module, which were not used in this study. The internal consistencies (Cronbach’s alpha) of the five items were 0.73 and 0.81 for the pre- and post-course surveys, respectively, indicating good internal consistencies of participants’ responses to the surveys.

### Course learning activities

Eight course learning activities (*Identity Grid, Role Grid, Mapping Your Goals Grid, Informational Interview Plan Grid, Putting Your Career Into Action, Social Identity Grid, Community of Practice Diagram, and Resilience Action Plan*) were developed to facilitate building deeper knowledge through application of module concepts (<u>S2 Table</u>). The grid-type activities consisted of multiple levels of open response questions to which learners responded, each building upon the last and progressing toward a goal, with video guidance and examples interspersed between stages (an example of *Putting Your Career Into Action* is included in <u>S1 Text</u>). The pedagogical design of these activities engaged learners in thinking through progressive steps to construct for themselves a process to advance and apply the skills they were learning. Learners needed to complete at least five activities to achieve course completion and receive a certificate. We chose six prompts (1. Obstacles on your pathway to success; 2. What I’m currently not doing 3. Why I’m currently not doing it 4. Strategies to overcome obstacles; 5. What will help you to achieve your goal? 6. What are the challenges/barriers you might encounter?) with detailed open responses from *Mapping Your Goals* and *Putting Your Career Into Action* from the eight course learning activities for qualitative analysis. The six prompts were chosen because they are open-ended questions and provided in-depth responses regarding learners’ awareness, knowledge, and ability to apply concepts than the single word or phrase entries in, for example, the *Identity Grid activity* (additional analysis of this and other activities will be the subject of a forthcoming manuscript, since they yield findings along different themes). We analyzed the responses as direct work products that speak to learners’ state of mind at that stage of the course and their own professional development.

## Data collection

The Institutional Review Board (IRB) of the researchers’ universities approved the study (IRB approval numbers is 5419X) and the participants who agreed to allow the information collected to be used for research were included in the study. Both surveys were embedded in the *Succeeding as a Postdoc* course on the edX platform and data were collected through Qualtrics. The pre-survey was administered during the first week of the course and the post-survey was administered in the sixth week of the course. Participants completed each survey in approximately 15 minutes. Course learning activities were embedded in *Succeeding as a Postdoc* and learners’ responses were collected and managed through PostgreSQL database as they progressed through each offering of the course.

## Data analysis

Data preparation and cleaning involved handling duplicates and missing values. If two response instances of the same individual were at the same level of completeness, the most recent one was retained. The percentage of missing values on the outcome variables ranged from 2% to 10% and that on predictor variables ranged from 8% to 11%. Little’s test [28] was first conducted using the R function *mcar_test* and the result revealed that the data were missing completely at random (MCAR). Multiple imputations were used to account for missingness using *m* = 5. Multiple imputation has been suggested to be advantageous over other alternative techniques (e.g., pairwise deletion, listwise deletion, single imputation) in handling missing values [29]. Prior to data analyses, we ensured that the assumptions for the repeated measures multivariate analysis of variance (MANOVA) and regressions had been met, which include independence, multivariate normality of dependent variables, linearity, homoscedasticity, and absence of multicollinearity [30].

A repeated measures MANOVA was conducted to determine if there were significant differences in participants’ self-perceptions of skills between the pre- and post-course surveys, our first research question. MANOVA was chosen instead of multiple ANOVAs or t-tests, because multiple tests would have increased the risk of committing Type I errors [30]. To answer the second research question, hierarchical linear regressions were conducted with gender, ethnicity, discipline, and country of origin entered as predictors and skills at the post-course survey entered as outcome variables. Skills in the pre-course survey were entered in the first step as the covariate. All the predictors were dummy coded. Ethnicity was coded as 1 for majority (i.e., White/Caucasian and Asian or Asian American) and 0 for URM (i.e., Alaska Native or Native American, Black or African American, Hispanic or Latino/Latina/Latinx, Middle Eastern or Northern African, Pacific Islander, and Multiracial groups). Gender was coded 1 for female and 0 for male, since we did not have sufficient number of non-binary participants. For discipline data, two dummy variables, discipline (biological/medical) and discipline (physical) were created with 0 being the reference variable and referring to the discipline of humanities/ social sciences, and 1 referring to biological/medical sciences and physical/engineering/computer sciences, respectively. Based on the distribution of countries by Human Development Index (HDI) [31], the variable of Country was dummy coded into the variable of country (highHDI), with 1 referring to very high and high HDI countries, and 0 referring to the medium and low HDI countries. The regression model is:

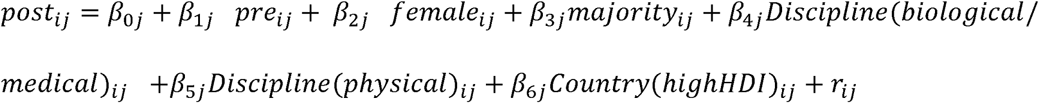

where j refers to each of the five skills (career transition, career planning, collaborative research, resilience, and self-reflection).

For the third research question regarding learners’ reflections on skill development, we conducted a thematic analysis [32] using Nvivo 12 [33] to examine learners’ responses to the six prompts of course learning activities. Coding occurred in two intentional phases. In the first phase of analysis, one researcher read all the open-ended responses. Open coding was adopted to locate and identify codes based on each prompt. After the code identification and exploration phase, coded information was reread to identify underlying connections between them and codes were categorized into themes and patterns. Several approaches were adopted to ensure the quality of data analysis and trustworthiness of findings. First, identified codes and themes were refined iteratively through dynamic discussions with team members. Second, 15% of the data was re-coded after three weeks to check diachronic reliability [34]. We obtained a reliability coefficient of 0.91, indicating the consistency and stability of the data analysis and interpretation. It is also important to note that the study team member who did the primary coding is a postdoc. The intersecting identity between the researcher and the participants contributed to the understanding of data and interpretation of findings. The remaining team members who contributed through discussion and alternative interpretations have extensive experience with research training at pre- and postdoc levels, and qualitative research methods.

## Results

### Self-perceptions of skill change

Results from the repeated measures MANOVA revealed that the omnibus Wilks’s lambda was statistically significant for the main effect of time, indicating that the combined dependent variables differed, on average, between pre-course and post-course surveys, Wilk’s = .623, *F*(5, 209) = 25.28, *P* < 0.001. From the univariate *F* analysis (Table 2), there were statistically significant increases over time for each measure. Learners self-reported skills in career transition, *F*(1, 213) = 25.23, *P* < 0.001, generalized *η*^2^ = .07, career planning, *F*(1, 213) = 44.97, *P* < 0.001, generalized *η*^2^ = .12, collaborative research, *F*(1, 213) = 18.46, *P* < 0.001, generalized *η*^2^ = .05, resilience, *F*(1, 213) = 29.04, *P* < 0.001, generalized ^2^ = .09, and self-reflection, *F*(1, 213) = 21.25, *P* < 0.001, generalized *η*^2^ = .07, all improved significantly after they took *Succeeding as a Postdoc*. Effect size is interpreted as small when *η*^2^ = 0.01, medium when *η*^2^ = 0.06, and large when *η*^2^ = 0.14 [35]. According to Cohen’s [35] benchmark, all the effect sizes ranged from medium to large, with the effect on career planning and resilience being the largest and the effect on collaborative research being the smallest.

**Table 2.**
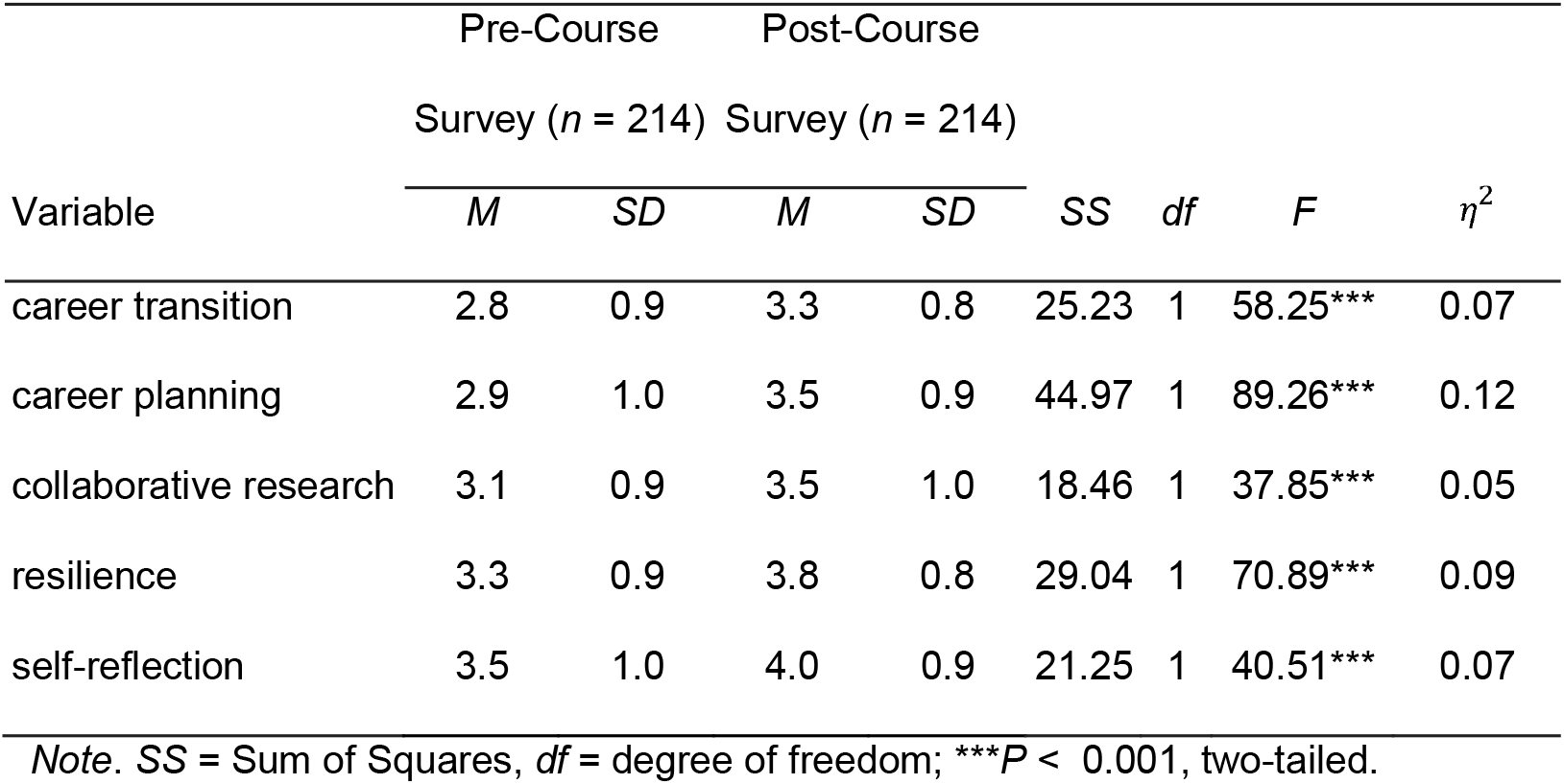
Means and Standard Deviations of Self-Reported Skills on the Pre- and the Post-Course Surveys, and the Follow-up Repeated Measures ANOVA Results. Data were from *The Postdoc Academy: Succeeding as a Postdoc* from February 2020 through January 2022 (pre-course survey, *n* = 214, post-course survey, *n* = 214).

|  | Pre-Course |  | Post-Course |  |  |  |  |  |
| --- | --- | --- | --- | --- | --- | --- | --- | --- |
|  | Survey (*n* = 214) |  | Survey (*n* = 214) |  |  |  |  |  |
| Variable | *M* | *SD* | *M* | *SD* | *SS* | *df* | *F* | $\eta^2$ |
| career transition | 2.8 | 0.9 | 3.3 | 0.8 | 25.23 | 1 | 58.25*** | 0.07 |
| career planning | 2.9 | 1.0 | 3.5 | 0.9 | 44.97 | 1 | 89.26*** | 0.12 |
| collaborative research | 3.1 | 0.9 | 3.5 | 1.0 | 18.46 | 1 | 37.85*** | 0.05 |
| resilience | 3.3 | 0.9 | 3.8 | 0.8 | 29.04 | 1 | 70.89*** | 0.09 |
| self-reflection | 3.5 | 1.0 | 4.0 | 0.9 | 21.25 | 1 | 40.51*** | 0.07 |

Note. SS = Sum of Squares, df = degree of freedom; \*\*\* $P < 0.001$ , two-tailed.

### Differences of skill changes by ethnicity, gender, discipline and country

Results from hierarchical linear regressions revealed that ethnicity had a significant influence on career planning (*t* = - 2.23, *P* < 0.05), resilience (*t* = - 3.87, P < 0.001), and self-reflection (*t* = - 2.09, *P* < 0.05) after controlling for differences in baseline skills on the pre-course survey. The unstandardized regression coefficient (B) indicated that URM learners reported 0.30, 0.47, 0.24 greater in their self-perceptions of career planning, resilience, and self-reflection, respectively, as compared to their majority counterparts. This result suggested that URM learners had greater gains in their self-perceptions of skills in the three aspects. However, no statistically significant differences were identified in perceived skill development as a function of gender, discipline, and country of origin (Table 3). In other words, although learners perceived that their skills improved after taking *Succeeding as a Postdoc*, the amount of skill development was not different between males and females, and among people from different disciplines and country backgrounds.

**Table 3.**
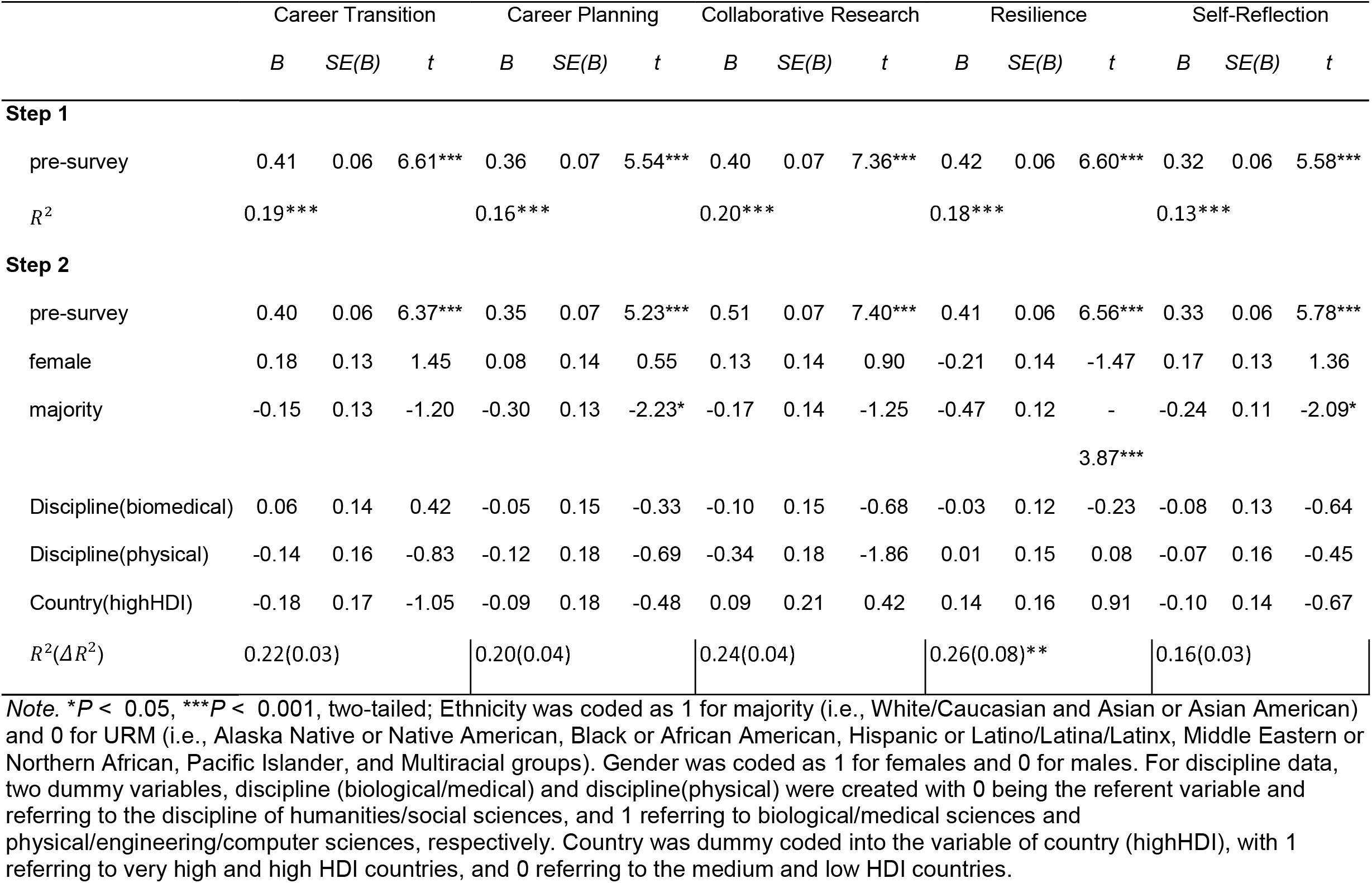
Pooled Hierarchical Regressions for Gender, Ethnicity, Discipline and Country of Origin Predicting Skill Development. Data were from *The Postdoc Academy: Succeeding as a Postdoc* from February 2020 through January 2022 (*n* = 214).

For this section and those that follow, we use “postdoc” to associate our findings with the large majority of learners since we demonstrate that no differences occur in the direction and significance of the estimates between the total data set and the sub-set that is only postdocs (see <u>S3</u> and <u>S4</u> Tables for the results with the sub-set of only postdocs).

### Reflections on learning from direct work product

Three themes were identified and the top (most frequent) four codes for each prompt between majority and URM learners were also compared (Fig 2) because significant differences in skill changes were found as a function of ethnicity. The top four codes were very similar, although the frequency order among the four differed between URM and majority postdocs.

**Fig 2.**
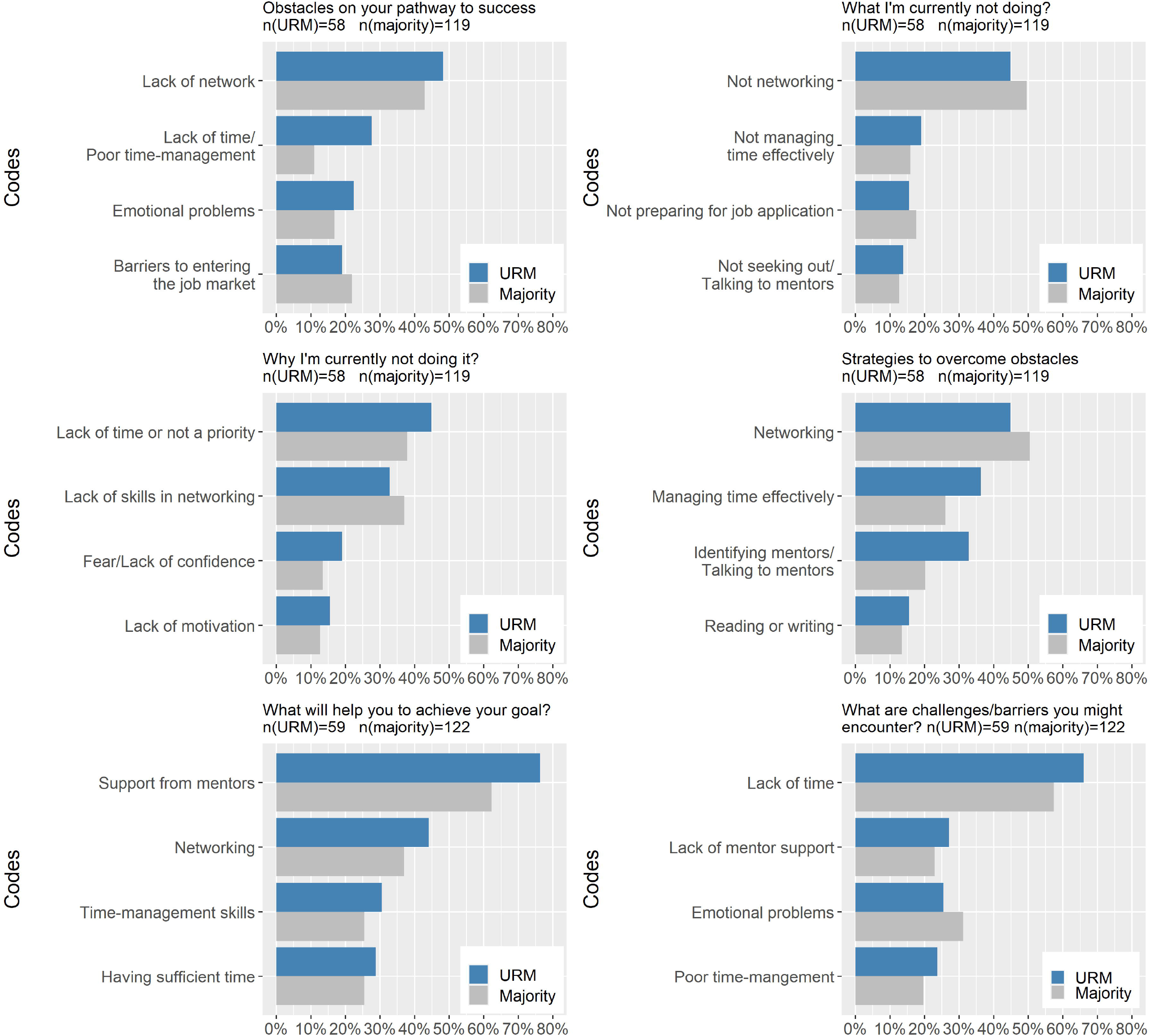
The Top Four Most Frequent Codes of Each Prompt for Majority and URM Learners. Data were from responses to the six open-ended prompts of learning activities (see above) from *The Postdoc Academy: Succeeding as a Postdoc* from February 2020 through January 2022.

### Networking

One of the most salient reflections among postdocs was that they perceived networking as necessary components for their career success. “Not networking” was one of the strong and consistent codes that emerged from participants’ responses to the prompt “What are you not currently doing”. Approximately half of the participants (45% of URM, *n* = 58, and 50% of majority, *n* = 119) mentioned that they neither “build their professional network with potential collaborators” nor “proactively reach out to other postdocs”.

> *“Outside of people I directly work/collaborate with for my research, my network is basically non-existent. I don’t do a good job at conferences introducing myself to people in my field and actually laying the foundation for my network”*.

However, postdocs realized that networking was a contributing factor to the success of their career as they engaged with the course, and it was among the top strategies that learners said they would use to overcome obstacles on their pathway to success. One postdoc made a plan to seek networking opportunities, saying:

> *“Over the next week, [I] will actively seek out and connect with individuals in my network who are professionals in research management or who have connections in the profession*.*”*

Participants noted that networking opportunities are necessary for successful career transition and planning and play a vital role in establishing research collaborations. One postdoc planned to improve networking by engaging with PIs and attending conferences and believed that s/he would build “possible collaboration with other colleagues”. In addition, connecting to “people who have gone through similar experiences” would help them “relieve stress, build confidence and develop resilience”.

### Perspectives on mentor support

Around one quarter of postdocs (27% of URM, *n* = 59, and 23% of majority, *n* = 122) perceived a lack of mentor support as a (potential) barrier to the achievement of their career goal, as evidenced in their responses to the prompt “What are the challenges/barriers you might encounter”. Despite that awareness, participants stated that they were not “proactively seeking out mentors” or “talking to existing mentors” due to the devotion of their time to other commitments such as research.

Postdocs reflected on the importance of mentorship while engaging with the course. Mentor support was the most salient theme emerging from their responses to the prompt “What will help you to achieve the goal” and a majority of learners believed that consistent and effective mentorship would bring intellectual or social benefits to their career success by offering “feedback and guidance” in writing manuscripts and grant proposals and providing “network opportunities and accessibility to resources”. This finding is prominent for URM postdocs, which was endorsed by 76% (*n* = 59) of URM postdocs as compared to 62% (*n* = 122) of majority postdocs. Thus, postdocs noted that they would either “identify a mentor” or “ask current mentors” for connections or information. One URM postdoc stated:

> *“[I would] [a]sk mentors for names of people to connect with or to actually connect me with them, try to start with someone I already know, talk to mentors about doing informational interviews so that I’m more likely to do it if I’ve talked to them about it*.*”*

### Challenges in engendering changes

Challenges are inevitable in any effort to engender and support change. Challenges that are salient in postdocs’ responses are a tension between a job and a career and fear of uncertainties.

#### The tension between fulfilling current job requirements and exploring a future career

Postdocs comprise a chronically overworked population, who enter needing to complete prior research, fulfill the obligations of current jobs, and prepare for their future careers [36]. A tension between a current job and a future career is a strong and consistent theme in our learners’ responses. Insufficient time and demanding workload were perceived as the main sources of their resistance to engaging in career preparation such as career-planning or collaboration-building behaviors, including conducting informational interviews, establishing social connections or seeking professional development opportunities. This theme is also more salient for URM learners. For example, over a quarter (28%, *n* = 58) URM postdocs perceived lack of time or time-management skills as a major obstacle on their way to success compared to only 11% (*n* = 119) for majority learners. Likewise, a majority of minority URM (66%, *n* = 59) believed that lack of time was the biggest challenge or barrier they might encounter in realizing their goals, which was endorsed by 57% (*n* = 122) of majority postdocs. In addition, 45% (*n* = 58) of URM and 38% (*n* = 119) of majority postdocs perceived “lack of time or not a priority” as the major reason that they were not seeking professional development opportunities or establishing networking. One postdoc commented:

> *“I’ve been heavily side-tracked with work and personal stuff that I often forget to contact others that I’ve met at conferences to network*.*”*

Faced with competing obligations, postdocs tended to prioritize their time to work on research-related tasks such as experiments, manuscripts and grants. One learner expressed this perspective by stating:

> *“Lack of time or better use of the time. I have been focused on writing manuscripts and performing experiments…So, the real challenge is considering this task as ‘minor’*.*”*

#### Fear of uncertainties

Learning activity responses revealed that postdocs were experiencing a multitude of negative emotions: imposter phenomenon, anxiety, stress, and low self-efficacy, which functioned as significant barriers to their proactive “help-seeking” or “reaching out” behaviors. Some also claimed that it was their “shy and intimidated” disposition that explained their hesitance to change. These negative emotions, coupled with the introverted disposition, induced a sense of uncertainty and fear, which would hold them back in networking or seeking collaboration opportunities. One learner listed their fears and hesitations:

> *“I’m afraid of being rejected when trying to interact with people. Also, I’m afraid of not understand[ing] when talking to people in English because sometimes they speak really fast. Additionally, I’m afraid of not being knowledgeable when talking to peers in conferences and I don’t want to look like a fool or stupid. And on top of all this I HATE SMALL TALK and I feel that is what I need to start conversations with new people*.*”*

## Discussion

We examined the changes of postdocs’ self-reported skills in career planning, transition, collaborative research, resilience and self-reflection as they engaged with a professional development course, *Postdoc Academy: Succeeding as a Postdoc*. We found that postdocs’ perceptions of their skills improved significantly upon completion of the course. Also, URM postdocs had greater gains in their self-perceived career planning, collaborative research and resilience, relative to majority ethnic postdocs. Qualitative analysis of course learning activities found that postdocs perceived networking and mentorship as support of their career progression, and revealed a tension between a current job and a future career and fear of uncertainties as significant barriers to improvement.

All five self-reported skills improved significantly, providing evidence of the effectiveness of *Postdoc Academy: Succeeding as a Postdoc* in supporting postdocs’ skill development in career transition, career planning, collaborative research, resilience and self-reflection, which are necessary for their success in and beyond a postdoc appointment. We hypothesize that the growth might be attributed to the inclusive and active-learning pedagogy, which created an accessible professional development environment for postdocs. Also, we believe videos that share stories and perspectives of diverse postdocs invite diverse learners to engage. We know from previous work that integrating self-reflection prompts and interactive activities create opportunities for participants to apply module concepts and directly facilitate their learning [5]. Notably, the course had larger effect sizes on the advancement of skills of career planning and resilience whereas had relatively smaller effect sizes on the growth of the skill of collaborative research, entirely consistent with the fewer number and depth of activities and course content related to collaborative research.

We found URM postdocs had greater gains in their self-reported development of skills in career planning, resilience and self-reflection as compared to their majority peers, suggesting that this course brings greater benefits to URM postdocs relative to majority postdocs. One explanation of this finding is that URM postdocs had lower level skills in the three aspects prior to entering the course, which might be caused by insufficient mentoring they received [17]. This provides support that our online, asynchronous course, *Succeeding as a Postdoc* can fill perceived needs in professional development and mentoring and is particularly beneficial for some URM postdocs.

We found postdocs perceived networking as key contributing factor to their skill advancement in preparation for job applications, identification of career and professional opportunities, and establishment of research collaboration and resilience. Personal and professional networks are vital for postdocs because many experience a sense of isolation and detachment, which are exacerbated by remote working opportunities and the COVID-19 pandemic [37]. This finding has been confirmed by the existing literature, which suggests that it is imperative for postdocs to establish a communication plan to expand their network or maintain connections with their local and disciplinary community [38].

We found that postdocs perceived mentor support as a significant factor influencing their career planning, transition, and collaborative research, and this perception was stronger for URM postdocs. This finding is consistent with previous research, which found that perceived mentor support during the postdoctoral period was a significant predictor of postdoc career plans [26], the odds of securing a permanent position, and future academic success [39]. McConnell et al. [26] further noted that mentor support was particularly beneficial for females and URM postdocs in their “pursuit of research-intensive academic careers”(p. 9). Our findings can be interpreted as a readily accessible, pedagogically inclusive and interactive means for URM postdocs to advance their skills, thus reducing the mentoring gap.

Our analysis found that the most significant challenges to postdocs’ applying their skills to advance career development are tensions among multiple current obligations and concern of future uncertainties. URM postdocs struggled with balancing multiple responsibilities and expressed a lack of confidence generally in skills necessary to achieve their careers, likely due to imposter phenomena and lack of belonging and identity [15, 40]. Hence *Succeeding as a Postdoc* provides some of the necessary professional development learning and practice to build skills and confidence, especially important for URM postdocs. We also found that postdocs were experiencing multiple negative emotions such as stress, anxiety, sadness and worry, which may be due to the nature of their positions (impermanence of their employment, heavy workload of current jobs, and insecurities of future career) and also the impact of the COVID-19 pandemic [41]. These negative emotions produced a sense of uncertainty, which were perceived as barriers to their cognitive or behavioral changes. Handled inappropriately, these emotions can lead to severe mental health issues and have a negative impact on academic achievement and professional success [21]. In the third module, *Developing Resilience*, we engage postdocs in learning, reflecting on, and applying resilience skills, and provided more information about resources and support services for emotional support or mental and psychological well-being.

## Limitations and future research

The current study is subject to limitations. First among them is that our participants comprised a self-selected and engaged subgroup of postdocs who completed the course learning activities and both the pre- and post-course surveys, which may pose an issue for generalizability and transferability of our findings. We do note, however, that we observed a high degree of homophily between the demographic distribution of our sample and the national postdoc population, indicating at least that such self-selection doesn’t shift the demographic representation. Second, the quantitative part of the study used a single-group pretest-posttest design, which may have threats to internal validity, including maturation and statistical regression (i.e., regression toward the mean). Third, the skills we measured were learners’ self-reported skills rather than their actual skills. Lastly, we found that URM postdocs tended to have more gains in their self-perceptions of skill development but we can only use prior studies and speculation as to why. Focus groups and interviews will be conducted in the future to obtain qualitative data to make the study more explanatory.

## Acknowledgements

We thank the following individuals for their expertise and assistance throughout all aspects of our study and for their help in improving the manuscript: Anne-Sophie Bohrer, Olivia Chesniak, Noah Green, Robin Greenler, Jessica Maher, Antonio Nunez, and Celine Young.

## Supporting information

**<u>S1 Table</u>. Demographic Information of Survey Respondents and National Postdoc Population**

**<u>S2 Table</u>. Brief Description of the Learning Activities in *Succeeding as a Postdoc***

**<u>S1 Text</u>. Description of *Putting Your Career Into Action* Learning Activity**

**<u>S3 Table</u>. Means and Standard Deviations of Self-Reported Skills on the Pre- and the Post-Course Surveys, and the Follow-up Repeated Measures ANOVA Results with the Sample of Postdocs Only**. Data were from *Succeeding as a Postdoc* from February 2020 through January 2022 (*n* = 178).

**<u>S4 Table</u>. Pooled Hierarchical Regression Analysis Summary for Gender, Ethnicity, Discipline and Country of Origin Predicting Skill Development with the Sample of Postdocs Only**. Data were from *Succeeding as a Postdoc* from February 2020 through January 2022 (*n* = 178).

## References

1. Feldon DF, Litson K, Jeong S, Blaney JM, Kang J, Miller C, et al. Postdocs’ lab engagement predicts trajectories of PhD students’ skill development. Proceedings of the National Academy of Sciences. 2019 Oct 15;116(42):20910-6.

2. Cavanaugh NA. Institutional support, programs, and policies for postdoctoral training. In The Postdoc Landscape 2018 Jan 1 (pp. 49–68). Academic Press.

3. Davis G. Doctors without orders: Highlights of the Sigma Xi postdoc survey. American Scientist. 2005 May 1;93(3):S1-.

4. Bessudnov A, Guardiancich I, Marimon R. A statistical evaluation of the effects of a structured postdoctoral programme. Studies in Higher Education. 2015 Oct 21;40(9):1588–604.

5. Hokanson SC, Grannan S, Greenler R, Gillian-Daniel DL, Campa H, Goldberg BB. A study of synchronous, online professional development workshops for graduate students and postdocs reveals the value of reflection and community building. Innovative Higher Education. 2019 Oct;44(5):385–98.

6. Ng TW, Eby LT, Sorensen KL, Feldman DC. Predictors of objective and subjective career success: A meta-analysis. Personnel psychology. 2005 Jun;58(2):367–408.

7. Lenzi RN, Korn SJ, Wallace M, Desmond NL, Labosky PA. The NIH “BEST” programs: Institutional programs, the program evaluation, and early data. The FASEB Journal. 2020 Mar;34(3):3570–82.

8. Layton RL, Solberg VS, Jahangir AE, Hall JD, Ponder CA, Micoli KJ, et al. Career planning courses increase career readiness of graduate and postdoctoral trainees. F1000Research. 2020;9.

9. NSF. Science and engineering indicators. 2016 [online] Available at https://www.nsf.gov/statistics/2016/nsb20161/uploads/1/nsb20161.pdf (p. 316, Section 3)

10. Phillips RA. Encouraging a more enterprising researcher: the implementation of an integrated training programme of enterprise for Ph. D. and postdoctoral researchers. Research in Post-Compulsory Education. 2010 Sep 1;15(3):289–99.

11. Hokanson SC, Goldberg BB. Proactive postdoc mentoring. In The Postdoc Landscape 2018 Jan 1 (pp. 91–120). Academic Press.

12. Subramanian S, Hutchins JA, Lundsteen N. Bridging the gap: increasing collaboration between research mentors and career development educators for PhD and postdoctoral training success. Molecular Biology of the Cell. 2022 Feb 1;33(2):pe1.

13. Lara LI, Daniel L, Chalkley R, editors. BEST: Implementing career development activities for biomedical research trainees. Academic Press; 2020 Feb 18.

14. Lambert WM, Wells MT, Cipriano MF, Sneva JN, Morris JA, Golightly LM. Research Culture: career choices of underrepresented and female postdocs in the biomedical sciences. Elife. 2020 Jan 3;9:e48774.

15. Yadav A, Seals C. Taking the next step: supporting postdocs to develop an independent path in academia. International Journal of STEM Education. 2019 Dec;6(1):1–1.

16. NIH. When it comes to applying for NIH grants and scholarships, who qualifies as an underrepresented person? 2021, January 21. URL https://www.niaid.nih.gov/grants-contracts/underrepresented-person-definition#:~:text=NIH%20considers%20the%20following%20groups,Individuals%20with%20disabilities

17. Beech BM, Calles-Escandon J, Hairston KG, Langdon MS, Latham-Sadler BA, Bell RA. Mentoring programs for underrepresented minority faculty in academic medical centers: a systematic review of the literature. Academic medicine: journal of the Association of American Medical Colleges. 2013 Apr;88(4).

18. Hariharan J. I felt lost in a new academic culture. Then I learned about the hidden curriculum. Science. 2019.

19. of Sciences NA, of Engineering NA. The Postdoctoral Experience Revisited.

20. NPA Core Competencies [WWW Document], n.d., URL https://www.nationalpostdoc.org/page/CoreCompetencies (Accessed 12 March 2022).

21. Gloria CT, Steinhardt MA. Relationships among positive emotions, coping, resilience and mental health. Stress and Health. 2016 Apr;32(2):145–56.

22. Wright MO, Masten AS, Narayan AJ. Resilience processes in development: Four waves of research on positive adaptation in the context of adversity. InHandbook of resilience in children 2013 (pp. 15-37). Springer, Boston, MA.

23. Scaffidi AK, Berman JE. A positive postdoctoral experience is related to quality supervision and career mentoring, collaborations, networking and a nurturing research environment. Higher Education. 2011 Dec;62(6):685–98.

24. Small G. The postdoc dilemma. Nature. 2012;483(7388):235.

25. Hara N, Solomon P, Kim SL, Sonnenwald DH. An emerging view of scientific collaboration: Scientists’ perspectives on collaboration and factors that impact collaboration. Journal of the American Society for Information science and Technology. 2003 Aug;54(10):952–65.

26. Mertens DM. Research and evaluation in education and psychology: Integrating diversity with quantitative, qualitative, and mixed methods. Sage publications; 2019 Feb 4.

27. McConnell SC, Westerman EL, Pierre JF, Heckler EJ, Schwartz NB. United States National Postdoc Survey results and the interaction of gender, career choice and mentor impact. Elife. 2018 Dec 18;7:e40189.

28. Chesniak OM, Drane D, Young C, Hokanson SC, Goldberg BB. Theory of change models deepen online learning evaluation. Evaluation and Program Planning. 2021 Oct 1;88:101945..

29. Little RJ. A test of missing completely at random for multivariate data with missing values. Journal of the American statistical Association. 1988 Dec 1;83(404):1198–202.

30. Yadav ML, Roychoudhury B. Handling missing values: A study of popular imputation packages in R. Knowledge-Based Systems. 2018 Nov 15;160:104–18.

31. Hahs-Vaughn DL. Applied multivariate statistical concepts. Routledge; 2016 Dec 1.

32. United Nations Development Programme. 2020 human development report. Retrieved from https://hdr.undp.org/en/composite/HDI.

33. Douglas E. Qualitative analysis: practice and innovation. Taylor & Francis; 2002.

34. QSR International Pty Ltd. (2020) NVivo (released in March 2020), https://www.qsrinternational.com/nvivo-qualitative-data-analysis-software/home

35. Ruan Z. Metacognitive awareness of EFL student writers in a Chinese ELT context. Language Awareness. 2014 Apr 3;23(1-2):76–91.

36. Cohen J. Statistical power analysis for the behavioral sciences. Routledge; 2013 May 13.

37. Small G. The postdoc dilemma. Nature. 2012;483(7388):235.

38. Fork ML, Anderson EC, Castellanos AA, Fischhoff IR, Matsler AM, Nieman CL, et al. Creating community: a peer□led, adaptable postdoc program to build transferable career skills and overcome isolation. Ecosphere. 2021 Oct;12(10):e03767.

39. Burgio KR, MacKenzie CM, Borrelle SB, Ernest SM, Gill JL, Ingeman KE, et al. Ten simple rules for a successful remote postdoc. PLoS Computational Biology. 2020 May 7;16(5):e1007809.

40. Liénard JF, Achakulvisut T, Acuna DE, David SV. Intellectual synthesis in mentorship determines success in academic careers. Nature communications. 2018 Nov 27;9(1):1–3.

41. Yadav, A., Soto, C., Smith, M., Marks, A., & Dixon, K. (2015, May). CIC AGEP Professorial Advancement Initiative. Paper presented at Understanding Interventions. San Diego, CA.

42. Woolston C. Pandemic darkens postdocs’ work and career hopes. Nature. 2020 Sep 8;585(7824):309–12.

