## Supplemental Table 1 for "A national professional development program fills mentoring gaps for postdoctoral researchers"

**S1 Table. Demographic information of survey respondents and national postdoc population**

| Variable | Category | Pre-Post Matched ($n$ = 215) | Pre-Survey ($n$ = 1151) | National Postdoc Population (McConnell et al., 2018) |
|---|---|---|---|---|
| Gender | Female | 146(67.9%) | 732(63.6%) | 4014(53.1%) |
| | Male | 49(22.8%) | 267(23.2%) | 3546(46.9%) |
| | Others | 2(0.9%) | 19(1.7%) | – |
| | Unknown | 18(8.4%) | 133(11.5%) | – |
| Ethnicity | White/Caucasian | 90(41.9%) | 525(45.6%) | 4674(60.3%) |
| | Asian or Asian American | 40(18.6%) | 221(19.2%) | 1924(24.8%) |
| | Hispanic or Latino/Latina/Latinx | 25(11.6%) | 100(8.7%) | 513(6.6%) |
| | Black or African American | 20(9.3%) | 47(4.1%) | 202(2.6%) |
| | Multiracial/More than one race | 14(6.5%) | 80(6.9%) | – |
| | Middle Eastern or Northern African | 4(1.9%) | 27(2.4%) | – |
| | Prefer to self-describe | 0 | 14(1.2) | – |
| | Unknown | 22(10.2%) | 137(11.9%) | – |
| Discipline | Biological/Medical Sciences | 120(55.8%) | 619(53.8%) | 5150(67.9%) |
| | Physical Sciences/Engineering /Computer Science | 30(13.9%) | 168(14.6%) | 1764(23.2%) |
| | Humanities and Social Sciences | 33(15.4%) | 173(15.0%) | 635(8.4%) |
| | Others | 12(5.6%) | 51(4.4%) | 36(0.5%) |
| | Unknown | 20(9.3%) | 140(12.2%) | – |
| Country | The United States | 92(42.8%) | 457(39.7%) | 3718(49.1%) |
| | Others | 100(46.5%) | 553(48.1%) | 3851(50.9) |
| | Unknown | 23(10.7%) | 141(12.2%) | – |

*Note.* Biological/Medical Sciences include biological and life sciences, medical sciences, agriculture and natural resource sciences, and earth, environmental, atmospheric, and ocean sciences; Physical Sciences/Engineering/Computer Sciences include computer, information, and technological sciences, engineering, mathematics and statistics, and physical sciences; Humanities/Social Sciences include Education, Humanities, Psychology, and Social, behavioral, and economic sciences.
