## Supplemental Text 1 for "A national professional development program fills mentoring gaps for postdoctoral researchers"

**S1 Text. Description of *Putting Your Career Into Action* learning activity**

The learning activity, Putting Your Career Into Action, was designed for learners to reflect and identify obstacles on their way to success and strategies to overcome those obstacles. First, learners are asked to identify potential obstacles on their pathway to success. Then, they pick a few of those obstacles that they can realistically build upon in the next few days and list what they are not doing. Third, learners are asked to reflect on why they are not doing those items and list why they are not doing those items. Lastly, learners reflect on how they are going to stop it and identify some strategies to overcome those obstacles. Below is a screenshot of this learning activity.

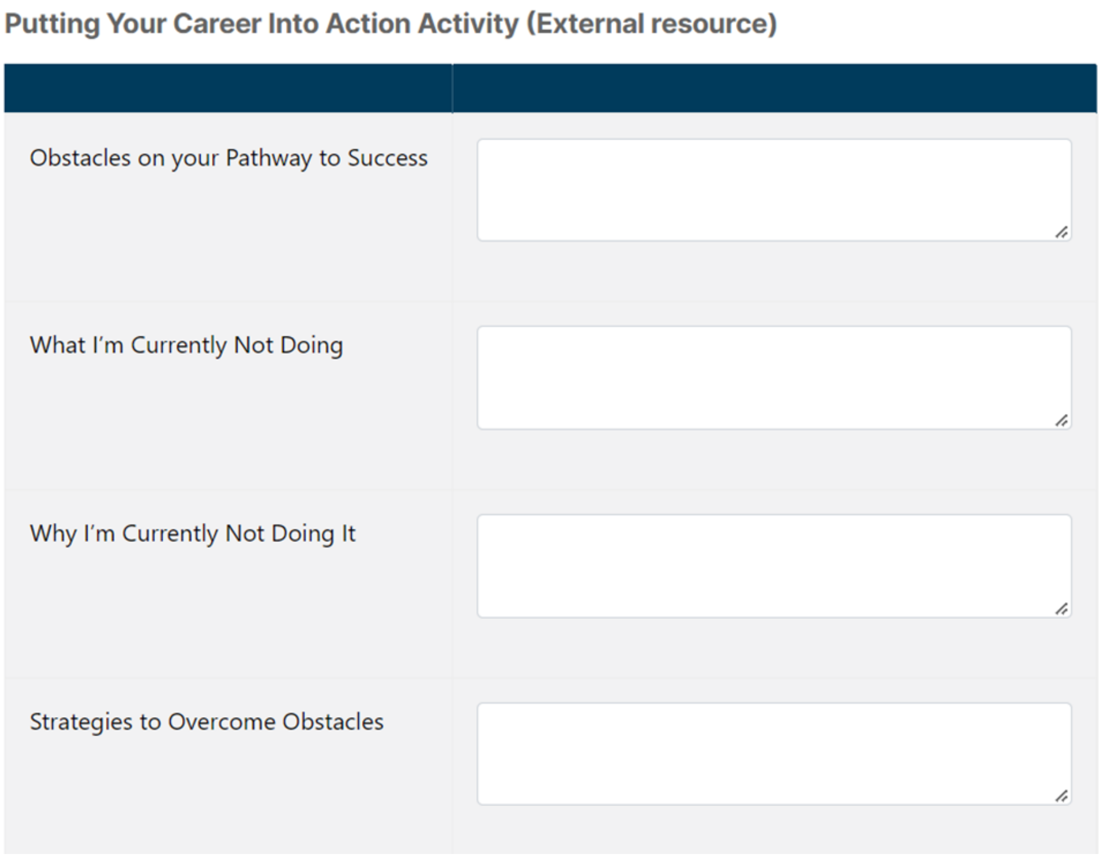
**Putting Your Career Into Action Activity (External resource)**

| |
|---|
| Obstacles on your Pathway to Success |
| What I'm Currently Not Doing |
| Why I'm Currently Not Doing It |
| Strategies to Overcome Obstacles |
