## Supplemental Table 2 for "A national professional development program fills mentoring gaps for postdoctoral researchers"

**S2 Table. Brief Description of the learning activities in *Succeeding as a Postdoc***

| Module | Leaning Activity | Instructions |
|---|---|---|
| Module 1 | Identity Grid Activity | Reflect on how you think about and define yourself. |
| | Role Grid Activity | Reflect on the transition you are going through and list a series of generic roles that postdocs have in their professional environment. |
| | Community of Practice Diagram | Create a diagram displaying your current research group and working relationships that exist across your group. |
| | Mapping Your Goals Grid Activity | Reflect on your goals and expectations for your postdoc and develop a timeline for your postdoc and the milestones that will lead to your success. |
| Module 2 | Informational Interview Plan Grid Activity | Plan for an informational interview. |
| | Putting Your Career into Action | Identify obstacles and strategies to overcome obstacles on your pathway to success. |
| Module 3 | Resilience Action Plan | Find an action plan for developing resilience. |
| Module 4 | Social Identity Grid Activity | Identify your social identities (gender, race, sex orientation, etc.) |
