## Supplemental Table 3 for "A national professional development program fills mentoring gaps for postdoctoral researchers"

**S5 Table. Means and standard deviations of self-reported skills on the pre- and the post-course surveys, and the follow-up repeated measures ANOVA results with the sample of postdocs only**. Data were from *The Postdoc Academy: Succeeding as a Postdoc* from February 2020 through January 2022 (pre-course survey, $n = 178$, post-course survey, $n = 178$).

| | Pre-Course Survey ($n$ = 178) | | Post-Course Survey ($n$ = 178) | | | | | |
|---|---|---|---|---|---|---|---|---|
| Variable | *M* | *SD* | *M* | *SD* | *SS* | *df* | *F* | $\eta^2$ |
| career transition | 2.8 | 0.9 | 3.3 | 0.8 | 23.18 | 1 | 47.37*** | 0.06 |
| career planning | 2.8 | 1 | 3.5 | 0.9 | 38.71 | 1 | 76.59*** | 0.13 |
| collaborative research | 3.1 | 0.9 | 3.5 | 1 | 14.17 | 1 | 28.18*** | 0.04 |
| resilience | 3.3 | 0.8 | 3.8 | 0.8 | 20.32 | 1 | 51.30*** | 0.08 |
| self-reflection | 3.5 | 1 | 4 | 0.8 | 18.85 | 1 | 37.27*** | 0.07 |

*Note*. *SS* = Sum of Squares, *df* = degree of freedom; ***$P < 0.001$, two-tailed.
