## Supplemental Table 4 for "A national professional development program fills mentoring gaps for postdoctoral researchers"

**S6 Table. Pooled hierarchical regression analysis summary for gender, ethnicity, discipline and country of origin predicting skill development with the sample of postdocs only.** Data were from *The Postdoc Academy: Succeeding as a Postdoc* from February 2020 through January 2022 (*n* = 178).

| | Career Transition | | | Career Planning | | | Collaborative Research | | | Resilience | | | Self-Reflection | | |
|---|---|---|---|---|---|---|---|---|---|---|---|---|---|---|---|
| | *B* | *SE(B)* | *t* | *B* | *SE(B)* | *t* | *B* | *SE(B)* | *t* | *B* | *SE(B)* | *t* | *B* | *SE(B)* | *t* |
| **Step 1** | | | | | | | | | | | | | | | |
| pre-survey | 0.34 | 0.08 | 4.34*** | 0.40 | 0.07 | 5.36*** | 0.49 | 0.08 | 5.89*** | 0.45 | 0.08 | 5.89*** | 0.34 | 0.07 | 5.17*** |
| $R^2$ | 0.13*** | | | 0.17*** | | | 0.2*** | | | 0.19*** | | | 0.15*** | | |
| **Step 2** | | | | | | | | | | | | | | | |
| pre-survey | 0.33 | 0.08 | 4.28*** | 0.40 | 0.07 | 5.43*** | 0.49 | 0.08 | 5.88*** | 0.43 | 0.07 | 5.88*** | 0.35 | 0.07 | 5.41*** |
| female | 0.31 | 0.16 | 1.88 | 0.19 | 0.17 | 1.07 | 0.13 | 0.19 | 0.70 | -0.16 | 0.17 | -0.97 | 0.15 | 0.15 | 0.98 |
| majority | -0.13 | 0.14 | -0.91 | -0.33 | 0.15 | -2.18* | -0.18 | 0.17 | -1.09 | -0.56 | 0.13 | -4.19*** | -0.34 | 0.13 | -2.52* |
| Discipline(biomedical) | 0.10 | 0.18 | 0.52 | 0.07 | 0.20 | 0.32 | -0.12 | 0.22 | -0.55 | -0.08 | 0.17 | -0.45 | 0.02 | 0.18 | 0.09 |
| Discipline(physical) | 0.10 | 0.21 | 0.49 | 0.08 | 0.24 | 0.32 | -0.09 | 0.25 | -0.37 | -0.09 | 0.20 | -0.43 | -0.09 | 0.21 | -0.43 |
| Country(highHDI) | -0.12 | 0.08 | -1.51 | -0.05 | 0.09 | -0.63 | 0.05 | 0.10 | 0.55 | 0.06 | 0.08 | 0.71 | -0.09 | 0.08 | -1.15 |
| $R^2(\Delta R^2)$ | 0.18(0.05) | | | 0.22(0.05） | | | 0.21(0.02) | | | 0.28(0.09)** | | | 0.22(0.07)* | | |
